# Social Instability is an Effective Chronic Stress Paradigm for both Male and Female Mice

**DOI:** 10.1101/550525

**Authors:** Christine N. Yohn, Sandra A. Ashamalla, Leshya Bokka, Mark M. Gergues, Alexander Garino, Benjamin A. Samuels

**Author notes:** Address correspondence to: Benjamin Adam Samuels, PhD, Assistant Professor, Behavioral & Systems Neuroscience, Department of Psychology, Busch Campus Psychology Building Room 215, Rutgers University - New Brunswick, 152 Frelinghuysen Road, Piscataway, NJ 08854.

## Abstract

Despite stress-associated disorders having a higher incidence rate in females, preclinical research mainly focuses on males. Chronic stress paradigms, such as chronic social defeat and chronic corticosterone administration, were mainly designed and validated in males and subsequent attempts to use these paradigms in females has demonstrated sex differences in the behavioral and HPA axis response to stress. Here, we developed a social stress paradigm, social instability stress (SIS), which exposes adult mice to unstable social hierarchies for 7 weeks. SIS effectively induces negative valence behaviors and hypothalamus-pituitary-adrenal (HPA) axis activation in both males and females. Importantly, while there were effects of estrous cycle on behavior, this variability did not impact the overall effects of SIS on behavior, suggesting estrous does not need to be tracked while utilizing SIS. Furthermore, the effects of SIS on negative valence behaviors were also reversed following chronic antidepressant treatment with fluoxetine (FLX) in both males and females. SIS also reduced adult hippocampal neurogenesis in female mice, while chronic FLX treatment increased adult hippocampal neurogenesis in both males and females. Overall, these data demonstrate that the SIS paradigm is an ethologically valid approach that effectively induces chronic stress in both adult male and adult female mice.

## INTRODUCTION

Several mood disorders, including anxiety and depression, are commonly thought to be precipitated and/or exacerbated by chronic exposure to stressful experiences. These stress-associated mood disorders occur at higher rates in women than in men. However, historically preclinical stress studies are notorious for only including males (1) and most chronic stress experimental paradigms are validated only using male rodents (2–4). One possible reason is that the female estrous cycle has well-documented effects on neural functions and behavior (5–8). However, it is unlikely that the estrous cycle contributes significantly more variability to female than male rodents (9,10). This realization, coupled with a mandate from the United States National Institutes of Health (NIH) to include both sexes in grant applications, has led to an increase in behavioral neuroscience studies that include both sexes (11). However, while some progress has been made, preclinical studies incorporating chronic stress still lag behind because many of the experimental paradigms were originally designed and optimized only for male rodents (1, 12–15).

One commonly used chronic stress paradigm in preclinical studies, unpredictable chronic mild stress (UCMS), differentially affects negative valence behaviors and HPA axis activation in male and female mice (16–19). Furthermore, UCMS has historically been plagued with reproducibility and validity issues (20–22). Another paradigm widely used in preclinical studies is chronic social defeat stress (CSDS), where rodents are subjected to larger more aggressive conspecifics (14). In males, CSDS results in activation of the hypothalamic-pituitary-adrenal (HPA) axis and increased negative valence behaviors (23, 24). However, standard CSDS protocols are not effective for stressing female rodents (25) unless variations that may infringe upon ethological validity such as activating the ventral medial hypothalamus of aggressors or applying male urine to females are made (26, 27).

In both males and females the HPA axis is activated by stress (28, 29), resulting in the release of the stress hormone cortisol (human) or corticosterone (rodents), and HPA axis dysregulation is found in many patients with mood disorders (30). Therefore, a distinct paradigm that effectively mimics chronic stress in male rodents is chronic administration of corticosterone (CORT). However, similar to CSDS, chronic CORT was developed in male rodents (13, 31, 32). In females, postpartum chronic administration of CORT increases immobility time within the forced swim test (FST) and decreases maternal behavior in female rats (33, 34). However, one recent study also suggests that chronic CORT is not as effective in producing increases in negative valence behaviors in females as in males (35).

Thus, there is currently no widely used chronic stress paradigm that is effective in both adult male and adult female rodents and many preclinical labs are either utilizing distinct paradigms for males and females or continuing to focus exclusively on males. Development of a single chronic stress paradigm that is effective in both sexes allowing direct comparisons and is ethologically valid is necessary for advancing our understanding of how chronic stress impacts neural function and behavior. Here, we describe the development of a social instability stress (SIS) paradigm that is effective in both adult male and adult female mice. The social instability paradigm described here involves exposure to unstable social hierarchical dynamics for several weeks and effectively induces negative valence behaviors, HPA axis activation, and neural changes associated with chronic stress such as altered adult hippocampal neurogenesis in both adult males and adult females of the widely used C57BL/6J strain. Furthermore, the negative valence behaviors induced by social instability in both adult males and adult females can be reversed by subsequent administration with the antidepressant fluoxetine, lending pharmacological validity to this chronic stress paradigm. Taken together, our results support the usage of social instability as an ethologically valid chronic stress paradigm that is effective for both adult male and adult female rodents.

## MATERIALS AND METHODS

### Mice

Adult 8 week old female and male C57BL/6J mice were purchased from Jackson laboratories and maintained on a 12L:12D schedule with *ad libitum* food and water. All testing was conducted in compliance with the NIH laboratory animal care guidelines and approved by Rutgers University Institutional Animal Care and Use Committee.

### Chronic Stress Paradigms

#### Chronic Corticosterone

Adult male and female C57BL/6J mice were randomly assigned to either vehicle (VEH) or corticosterone (CORT) treatment, with weights measured once per week during treatments. Corticosterone (35 ug/mL, equivalent to 5 mg/kg/day) was dissolved in 0.45% beta-cyclodextrin (Sigma) water and delivered ad libitum in opaque drinking bottles (David et al., 2009). VEH mice received 0.45% beta-cyclodextrin water ad libitum. Following 4 weeks of CORT or VEH treatment, mice received 3 weeks of fluoxetine (FLX) (18 mg/kg/day) or VEH (water) via oral gavage with CORT or VEH remaining in the drinking water throughout antidepressant treatment (timeline in Figure 1A). On behavioral testing days FLX or VEH was administrated after mice completed the behavioral tests to avoid acute effects. The following 4 groups emerged: VEH+VEH, VEH+FLX, CORT+VEH, and CORT+FLX.

**Figure 1.**
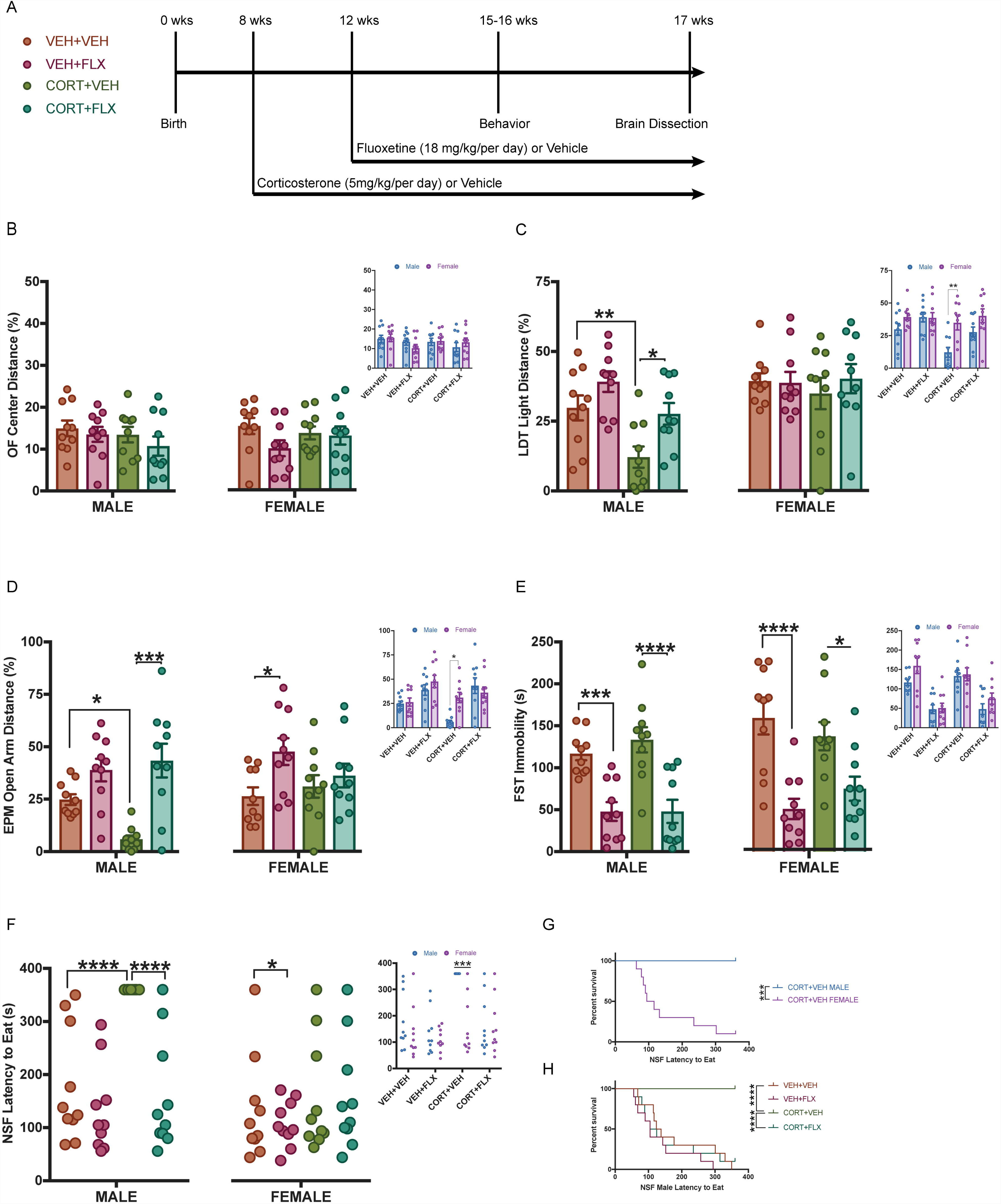
Behavioral Effects of Chronic Corticosterone Administration in Male and Female Mice. (A) Timeline of chronic CORT paradigm and keys for larger graphs. (B-E) Small panels represent 2×4 ANOVAs Bonferroni posthocs of Sex*x*Group (VEH+VEH, VEH+FLX, CORT+VEH, CORT+FLX) with sex differences observed in distance traveled in light (C small panel: *F*_(1,72)_=13.69, *p*<0.001, **CORT+VEH males *vs* CORT+VEH females *p*=0.001) and on open arms of EPM (D small panel: *F*_(1,72)_=9.97, *p<*0.001, *CORT+VEH males *vs* CORT+VEH females *p*=0.005). Larger (B-E) panels represent separate 2×2 ANOVAs Bonferroni posthocs (CORT*x*FLX) within each sex. Differences in males were observed in LDT (CORT: *F*_(1,36)_=13.53, *p*=0.0008; FLX: *F*_(1,36)_=9.88, *p*=0.0033; **CORT+VEH*vs*VEH+VEH *p*=0.0007 and *CORT+VEH*vs*CORT+FLX (*p*=0.0017), EPM open arm distance (FLX: *F*_(1,36)_=25.68, *p*<0.0001, INTERACTION: *F*_(1,36)_=5.29, *p*=0.027; *CORT+VEH*vs*VEH+VEH *p*=0.025 and **CORT+VEH*vs*CORT+FLX *p*=0.0042), and FST immobility (FLX: *F*_(1,36)_=39.38, *p*<0.00001; ***VEH+VEH*vs*VEH+FLX *p*<0.0001, ****CORT+VEH*vs*CORT+FLX *p*<0.0001). Smaller scatterplot (F) represents sex differences observed by Kaplan Meier survival analysis (G) ***CORT+VEH males *vs* CORT+VEH females *p*<0.001. Larger scatterplot (F) shows NSF data showing individual latency to eat values with Kaplan Meier survival analysis in males (H) ****CORT+VEH*vs*VEH+VEH *p*<0.0001; ****CORT+VEH*vs*CORT+FLX, *p*<0.0001). Additionally, differences in female detected in EPM open arm distance (D; FLX: *F*_(1,36)_=5.88, *p*=0.02; *VEH+VEH*vs*VEH+FLX *p*=0.018) and FST immobility (E; FLX: *F*_(1,35)_=28.37, *p*<0.0001, ****VEH+VEH*vs*VEH+FLX p<0.0001, *CORT+VEH*vs*CORT+FLX *p*=0.018). Scatterplot (F) represents feeding behavior differences form Kaplan Meier survival analysis between *VEH+VEH and VEH+FLX (*p*=0.033).

#### Social Instability Stress (SIS)

Adult male and female C57BL/6J mice were randomly assigned to either SIS or control (CNTRL, no stress). SIS mice experienced unstable social hierarchies, where social dynamics were changed twice a week for 7 weeks (adapted from 36–38). During the 7 weeks, a cage composition change would consist of an individual mouse being introduced to 3-4 novel mice of the same sex, with total mice per cage ranging from 3-5 mice. The rotation schedule was randomized to prevent mice from having an encounter with a recent cage mate. At the end of the 7 weeks, SIS mice remained housed with the mice of the last cage composition (adapted from 36). Male and female CNTRL mice had the same cage mates throughout the entire paradigm and were subjected to cage changes twice per week (adapted from 36, 39). Subsequent to SIS exposure, SIS and CNTRL mice were randomly assigned to receive either FLX (18 mg/kg/day) or VEH (water) via oral gavage (timeline in Figure 2A). The following 4 groups emerged: CNTRL+VEH, CNTRL+FLX, SIS+VEH, sand SIS+FLX.

**Figure 2.**
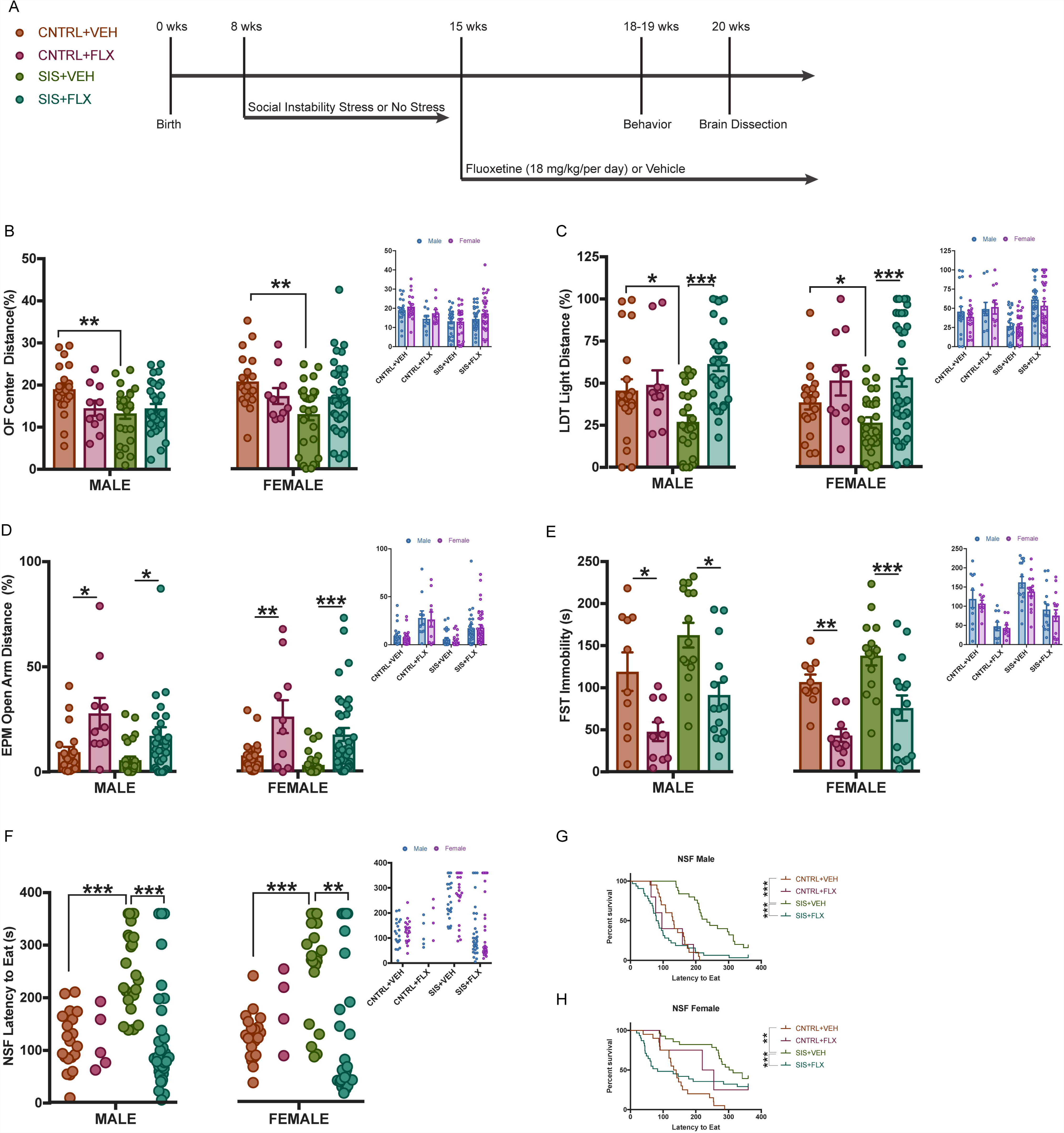
Behavioral Effects of Social Instability Stress (SIS) in Male and Female Mice. (A) Timeline of SIS paradigm and keys for larger graphs. (B-E) Small panels represent 2×4 ANOVAs Bonferroni posthocs of Sex*x*Groups (CNTRL+VEH, CNTRL+FLX, SIS+VEH, SIS+FLX) with no sex differences observed in any behavior. Larger (B-E) panels represent separate 2×2 ANOVAs Bonferroni posthocs (SIS*x*FLX) within each sex. Differences in males were observed in OF (SIS: *F*_(1,83)_=4.148, *p*=0.0449; interaction: *F*_(1,83)_=4.087, *p=*0.0464; **CNTRL+VEH*vs*SIS+VEH, *p*=0.0041) as well as in females; (SIS: *F*_(1,91)_=5.017, *p*=0.0276: SISxFLX: *F*_(1,91)_=4.643, *p=*0.0339; **CNTRL+VEH*vs*SIS+VEH, *p*=0.0019). LDT (CORT: *F*_(1,83)_=10.53, *p*=0.0017; FLX: *F*_(1,83)_=7.076, *p*=0.0094; **CNTRL+VEHvsSIS+VEH *p*=0.029 and ***SIS+VEHvsSIS+FLX (*p*<0.0001), EPM open arm distance (FLX: *F*_(1,36)_=25.68, *p*<0.0001, INTERACTION: *F*_(1,36)_=5.29, *p*=0.027; *CORT+VEHvsVEH+VEH *p*=0.025 and **CORT+VEH*vs*CORT+FLX *p*=0.0042), and FST immobility (FLX: *F*_(1,36)_=39.38, *p*<0.00001; ***VEH+VEH*vs*VEH+FLX *p*<0.0001, ****CORT+VEH*vs*CORT+FLX *p*<0.0001). Smaller scatterplot (F) represents sex differences observed by Kaplan Meier survival analysis (G) ***CORT+VEH males *vs* CORT+VEH females *p<*0.001. Larger scatterplot (F) shows NSF data showing individual latency to eat values with Kaplan Meier survival analysis in males (H) ****CORT+VEH*vs*VEH+VEH *p<*0.0001; ****CORT+VEH*vs*CORT+FLX, *p*<0.0001). Additionally, differences in female detected in EPM open arm distance (D; FLX: *F*_(1,36)_=5.88, *p*=0.02; *VEH+VEH*vs*VEH+FLX *p*=0.018) and FST immobility (E; FLX: *F*_(1,35)_=28.37, *p*<0.0001, ****VEH+VEH*vs*VEH+FLX *p*<0.0001, *CORT+VEH*vs*CORT+FLX *p*=0.018). Scatterplot (F) represents feeding behavior differences form Kaplan Meier survival analysis between *VEH+VEH and VEH+FLX *p*=0.033.

### Vaginal Lavage

Vaginal lavages were performed daily during the stress paradigms and two weeks prior to behavioral testing to ensure mice were cycling throughout all four stages of the estrous cycle regularly. After completing each behavioral test, vaginal smears were collected to assess the estrous state mice were in during the behavior test. Samples were collected via a pipette filled with ddH_2_O gently expelled and placed at the vaginal canal opening (without penetration). Samples were suctioned back into the pipette, placed on a microscope slide, and dried on a slide warmer before imaged with an EVOS FL Auto 2.0 microscope (Thermofisher Scientific) at 10x magnification (40, 41). Estrous phases were identified by the presence or absence of nucleated epithelial cells, cornified epithelial cells, and leukocytes (41, 42 Figure 3B).

**Figure 3.**
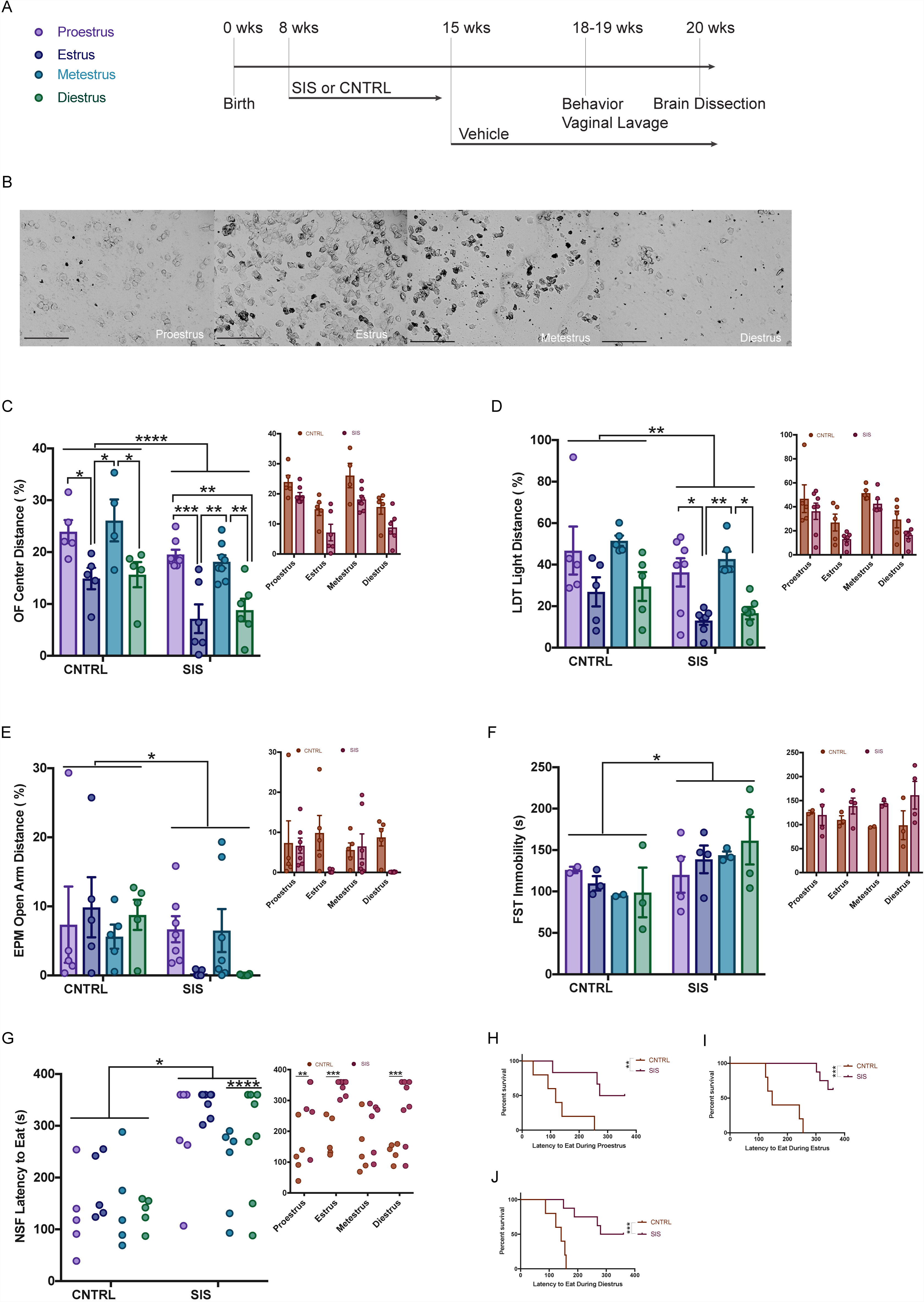
Impact of Estrous Cycle in Females on Behavior. Time line of experiment (A) and representative 10x images of vaginal smears (B). Larger panels (C-F) represent 2×4 ANOVAs (SIS*x*ESTROUS) in the OF (****SIS: *F*_(1,39)_=19.13, *p*<0.0001; ESTROUS: *F*_(3,39)_=45.16, *p*<0.001; ***SIS+ESTRUS*vs*SIS+PROESTRUS *p=*0.0003; **SIS+ESTRUS*vs*SIS+METESTRUS *p=*0.013; **SIS+DIESTRUS*vs*SIS+PROESTRUS *p=*0.0016; **SIS+DIESTRUS*vs*SIS+METESTRUS *p=*0.0094; ***CNTRL+ESTRUS*vs*CNTRL+PROESTRUS *p=*0.045; **CNTRL+ESTRUS*vs*CNTRL+METESTRUS *p=*0.013; *CNTRL+DIESTRUS*vs*CNTRL+PROESTRUS *p=*0.022; *CNTRL+DIESTRUS*vs*CNTRL+METESTRUS *p=*0.023), LDT (**SIS: *F*_(1, 39)_=7.68, *p=*0.0085; ESTROUS: *F*_(3,39)_=10.43, *p*<0.0001 Bonferroni corrected comparisons *SIS+ESTRUS*vs*SIS+PROESTRUS *p=*0.021; **SIS+ESTRUS*vs*SIS+METESTRUS *p=*0.003; *SIS+DIESTRUS*vs*SIS+METESTRUS *p=*0.011), EPM (*SIS: *F*_(1, 39)_=5.87, *p=*0.0196), and FST (*SIS: *F*_(1, 17)_=4.47, *p=*0.04). No differences within estrous state between SIS and CNTRL were observed smaller panels (C-F). Scatterplot (G) form Kaplan Meier survival analysis of latency to eat across estrous stages within the SIS group (χ^2^(3)=11.71, *p=*0.0084; ****SIS+ESTRUS*vs*SIS+METESTRUS *p*<0.0001). Comparison between SIS and CNTRL across estrous cycle (smaller panel G) represent differences in Kaplan Meier survival analysis of latency to eat across **SIS+PROESTRUS*vs*CNTRL+PROESTRUS *p=*0.0085 (H), ***SIS+ESTRUS*vs*CNTRL+ESTRUS *p=*0.0001 (I), ***SIS+DIESTRUS*vs*CNTRL+DIESTRUS *p=*0.0001 (J).

### Behavioral Testing

#### Open Field (OF)

Motor activity was quantified via infrared photobeams in Plexiglass open field boxes 43×43 cm^2^ (Kinder Scientific). As previously described (13), activity chambers were computer interfaced for data sampling at 100ms resolution. The computer software predefines grid lines that divide each OF chamber into center and periphery regions, with the center being a square 11cm from the wall. For our analyses we calculated percent distance traveled in the center ((center distance/total distance traveled)* 100).

#### Light Dark Test (LDT)

LDT was conducted in OF chambers with, a dark plastic box (opaque to visible light, but transparent to infrared light) covering 1/3 of the arena, inserted to separate the OF into light and dark compartments. The dark box contained an opening that allowed passage between the light and dark (13), with the light compartment brightly illuminated (1000 lux; Kinder Scientific). At the beginning of each 5-minute test, mice were placed in the dark compartment, with distance traveled in the light ((distance traveled in the light/total distance) *100) used for analyses.

#### Elevated Plus Maze (EPM)

The EPM test consisted of a plus-shaped apparatus with two open and two closed arms (side walls), elevated 2 feet from the floor. During the five-minute test, the mice were recorded from a video camera mounted on the ceiling above each EPM arena. EthoVision (Noldus) software was used to quantify the data, with distance traveled in open arms ((total open arm distance/total distance traveled)*100) used for analyses.

#### Forced Swim Test (FST)

A modified FST procedure suitable for mice was used (13), with individual cylinders (46×32×30 cm) filled with room-temperature water (25-26°C). Two sets of photobeams were mounted on opposite sides of the cylinder (Kinder Scientific) to record swimming behavior during the 6-minute test. Immobility was assessed during only the last 4-minutes of the test since mice habituate to the task during the initial 2-minutes.

#### Novelty Suppressed Feeding (NSF)

Mice were food deprived for 18 hours within their home cage, prior to being placed in the corner of a testing apparatus (50×50×20 cm) filled with 2 cm of corncob bedding, with a single food pellet attached to a white platform in the brightly illuminated center (1500 lux). The NSF test lasted 6 minutes with latency to eat (defined as the mouse sitting on its haunches and biting the pellet with the use of forepaws) recorded in seconds. Mice that timed out were assigned a latency of 360secs. Immediately after the test, mice were transferred to their home cages and given ad libitum access to food for 5 minutes. Latency to eat and amount of food consumed in home cage was measured as a control for feeding behavior observed in the NSF.

### Brain Collection, Sectioning, and Immunohistochemistry

#### Brain Collection and Sectioning

Following behavioral testing, brains were collected from all experimental mice. Mice were anesthetized with ketamine (80mg/kg) and perfused transcardially with PBS followed by 4% paraformaldehyde. Brains were collected, stored in 4% paraformaldehyde overnight at 4°C, and then switched to 30% sucrose 0.1% sodium azide (NaN_3_) in PBS solution until sectioned. Using a cryostat, the hippocampus was collected (43; Bregma −1.22 to −3.88) and mounted on SuperfrostPlus slides (Thermofisher Scientific) and stored at −20°C until immunostaining.

#### Immunohistochemistry

The effects of SIS and FLX on adult hippocampal neurogenesis were assessed across the entire hippocampus by counting 1 out of every 6 hippocampal sections. Slides were washed in 1% TritonX-100-PBS and then 3 PBS washes. Next, slides were incubated in citrate buffer for 30 minutes followed by 3 PBS washes. Slides were blocked for 1 hour in 10% normal goat serum (NGS) diluted in PBS before being incubated overnight at 4°C in either anti-rabbit Ki67 (1:500; Abcam, ab16667) or doublecortin anti-rabbit (1:500; Life technologies; 481200) diluted in 2%NGS-PBS. Following 18 hours of incubation, 3 PBS washes were performed and then 2 hours of incubation in CY-5 goat anti-rabbit (1:1000, Invitrogen, ThermoFisher Scientific, A10523) diluted in 2%NGS-PBS. Following 3 more PBS washes, slides were counterstained with DAPI (1:15000; ThermoFisher Scientific) for 15 minutes, with a final PBS wash before cover slipping using prolong diamond (ThermoFisher Scientific). High-resolution fluorescent images were taken using an EVOS FL Auto 2.0 microscope (Thermofisher Scientific) at 10x magnification for quantification of Ki67^+^ (Figure 5A) or DCX^+^ cells (Figure 5C) and counted across a total of 12 sections of hippocampus. Ki67^+^ cells were overlaid with DAPI for quantification purposes. Images were also taken at 40x magnification to subcategorize DCX^+^ cells according to their dendritic morphology: DCX^+^ cells with no tertiary dendritic processes and DCX^+^ cells with complex, tertiary dendrites (Figure 5E). The maturation index was defined as the ratio of DCX^+^ cells possessing tertiary dendrites over the total DCX^+^ cells.

### Blood Collection and Corticosterone ELISA

Mice were weighed to ensure that non-terminal blood collection was no more than 1% of the mouse’s body weight. Prior to SIS, all mice had baseline blood samples collected from the left retro-orbital sinus in accordance with IACUC guidelines. To measure corticosterone levels in response to a change in social dynamics (i.e. cage composition change), male and female SIS mice had blood collected from the right retro-orbital sinus approximately 40-45 minutes after cage change. CNTRL mice had blood collected from the right retro-orbital sinus 40-45 minutes after a cage change, to control for impact of exposure to a novel cage on plasma corticosterone levels. Blood was also collected 40-45 minutes following the EPM from the left retro-orbital sinus of SIS and CNTRL mice to measure plasma corticosterone levels in response to a negative valence behavior. For each of the three blood collections, blood was collected in micro centrifuge tubes coated with EDTA. Plasma was isolated from whole blood by centrifugation at 14000 rpm for 10 min at 4 °C, with supernatant collected and stored at - 80°C until assayed. Total yield of plasma per blood collection was between 25 to 40 µL. To assess differences in plasma corticosterone in response to SIS stress and EPM behavior, blood was analyzed from each time point (prior to SIS, SIS cage change, EPM) across 5 mice per sex per stress condition (male: CNTRL=5, SIS=5; female: CNTRL=5, SIS=5). Each sample (total of n=60 across all time points) was diluted 1:100 and assayed in triplicate according to the manufacturer’s protocol (Arbor Assays Corticosterone ELISA Kit).

### Statistical Analyses

All analyses were conducted using Graph Pad Prism 7. Sex differences for both CORT and SIS paradigms were analyzed using 2×4 analysis of variance (ANOVA) with follow-up Bonferroni post-hoc comparisons. Further analyses in the SIS paradigm used 2×2 ANOVAs within sex to compare stress (SISvsVEH) and FLX (FLXvsVEH) effects. Given that the NSF data fails to meet basic assumptions of normality, Kaplan Meier survival analysis (nonparametric test) was used to analyze sex differences in feeding behavior. Lastly, to analyze differences in exogeneous corticosterone levels across the three time points (prior to SIS, SIS cage change, and EPM exposure), time point, sex, and groups were used to conduct a 3×2×2 repeated measures ANOVA. These analyses of corticosterone levels were then followed up with separate 2×3 repeated measures ANOVAs within each sex.

## RESULTS

### Behavioral Sex Differences following Chronic CORT Administration

Chronic CORT administration is used to mimic chronic stress in male rodents (13, 31). However, there are inconsistencies in published reports about whether CORT administration is also effective in female rodents (33–35). Therefore, we chronically administered exogenous CORT (5mg/kg/day) or VEH to male and female C57BL/6J mice (timeline in Figure 1A). Following 4 weeks of CORT, mice received either FLX (18mg/kg) or VEH, resulting in four groups: VEH+VEH (male=10; female=10), VEH+FLX (male=10; female=10), CORT+VEH (male=10; female=10), and CORT+FLX (male=10; female=9). We examined CORT and FLX effects across these groups in negative valence behavioral measures known to be affected by chronic stress paradigms: open field (OF), light/dark test (LDT), elevated plus maze (EPM), novelty suppressed feeding (NSF), and forced swim test (FST) respectively.

First, separate 2×4 ANOVAs (sex *x* group) were used to investigate potential sex differences. Significant sex effects were found in LDT light distance traveled (*F*_(1,72)_=13.69, *p*<0.001; Figure 1C small panel) and EPM open arms (*F*_(1,72)_=9.97, *p*<0.001; Figure 1D small panel), with CORT+VEH males traveling less distance in the light (*p*=0.001) and in EPM open arms (*p*=0.005) than CORT+VEH females (Figure 1C-D small panels). Additionally, sex differences in NSF latency to eat were discovered by Kaplan Meier survival analyses (log-rank Mantel Cox test), with CORT+VEH males having a longer latency to eat than CORT+VEH females (χ^2^_(1)_=37.35, *p*<0.001; Figure 1G). No sex differences were observed between groups in the OF (*p*=0.97) and FST (*p*=0.17). Taken together, these data demonstrate that CORT administration more effectively induces negative valence behaviors in males than in females.

After examining sex differences, separate 2×2 ANOVAs were run to investigate CORT and FLX effects on negative valence behaviors within each sex. Within males significant CORT (*F*_(1,36)_=13.53, *p*=0.0008) and FLX (*F*_(1,36)_=9.88, *p*=0.0033) effects on LDT light distance traveled (Figure 1C) were found, with CORT+VEH males traveling less than VEH+VEH (*p*=0.0007) and CORT+FLX (*p*=0.0017) males (Figure 1C larger panel; Bonferroni corrected). In the EPM, significant FLX (*F*_(1,36)_=25.68, *p*<0.0001) and interaction (*F*_(1,36)_=5.29, *p*=0.027) effects were observed in open arm distance traveled, with CORT+VEH males traveling less in the open arms than VEH+VEH (*p*=0.025) and CORT+FLX (*p*=0.0042) males (Figure 1D larger panel; Bonferroni corrected). Additionally, in males a significant effect of FLX (*F*_(1,36)_=39.38, *p*<0.00001) was observed on FST immobility time, with CORT+FLX males being less immobile than CORT+VEH males (*p*<0.001, Bonferroni corrected; Figure 1E). VEH+FLX males were also less immobile than VEH+VEH males (*p*=0.0007; Bonferroni corrected; Figure 1E). Lastly, Kaplan Meier survival analysis revealed a significant difference between male groups (χ^2^_(3)_= 37, *p*<0.0001), with further analyses revealing CORT+VEH males had a longer latency to eat than VEH+VEH (χ^2^_(1)_= 37.58, *p*<0.0001; Bonferroni corrected) and CORT+FLX (χ^2^_(1)_= 15.03, *p*<0.0001; Bonferroni corrected; Figure 1F-H).

In females, separate 2×2 ANOVAs investigating the effects of CORT and FLX showed a significant FLX effect in the EPM (*F*_(1,36)_=5.88, *p*=0.02) and FST (*F*_(1,35)_=28.37, *p*<0.0001), with VEH+FLX females spending more time in the open arms (*p*=0.018, Bonferroni corrected; Figure 1D) and less time immobile (*p*<0.0001; Figure 1E) than VEH+VEH females. CORT+FLX females also spent less time immobile in the FST than CORT+VEH females (*p*=0.018; Figure 1E). Lastly, Kaplan Meier survival analysis revealed a significant difference between female groups (χ^2^_(3)_= 9.28, *p*=0.026), with further analyses revealing VEH+FLX females had a shorter latency to eat than VEH+VEH females (χ^2^_(1)_= 4.54, *p*=0.033; Bonferroni corrected; Figure 1F). Taken together, these data further demonstrate that while FLX has behavioral effects in both males and females, CORT administration only mimics the effects of chronic stress in males. These data support the findings of Mekiri and colleagues (35) and suggest that CORT administration is not a useful paradigm for studying stress effects in females.

### SIS is Effective in both Males and Females

Since chronic CORT appeared to only mimic the behavioral effects of chronic stress in males, we next developed a social instability paradigm that we reasoned would be stressful for both sexes. To this end, C57BL/6J mice were exposed to unstable same sex hierarchical conditions within their housing environment (Figure 2A). Specifically, SIS mice experience a change in cage dynamics every 3 days for 7 weeks, while CNTRL are housed with the same cage mates for the duration of the experiment. Adult male and adult female C57BL/6J mice were assigned to either SIS or CNTRL (no stress) housing conditions for 7 weeks, and then received 3 weeks of either VEH or FLX (18mg/kg daily) resulting in the following groups: CNTRL+VEH (male=20; female=20), CNTRL+FLX (male=10; female=10), SIS+VEH (male=25; female=28); SIS+FLX (male=33; female=35). To analyze whether sex differences existed within the four groups (CNTRL+VEH, CNTRL+FLX, SIS+VEH, and SIS+FLX), 2×4 ANOVAs (sex x group) were run for each behavior. All 2×4 ANOVAs analyses revealed no main effect of sex for any measure: distance traveled in OF center (*p*=0.11), LDT light distance (*p*=0.44), EPM open arm distance (*p*=0.65), FST immobility (*p*=0.17) (Figures 2B-F small panels). For NSF, Kaplan Meier survival curves also showed no sex differences across the four groups (*p*=0.33) (Figure 2F-H). These data suggest that SIS may be impacting males and females similarly. Thus, to determine the effectiveness of SIS exposure and subsequent FLX treatment in both males and females, 2×2 ANOVAs (SISxFLX) were run within each sex.

In males, a 2×2 ANOVA (SISxFLX) revealed significant SIS effect (*F*_(1,83)_= 4.148, *p*=0.0449) and interaction (*F*_(1,83)_= 4.087, *p*=0.0464) in OF center distance traveled, with SIS+VEH males traveling less distance than CNTRL+VEH males (*p*=0.0041; Figure 2B larger panel; Bonferroni corrected). Significant main effects of FLX (*F*_(1,83)_= 10.53, *p*=0.0017) and an interaction (*F*_(1,83)_= 7.076, *p*=0.0094) were found in LDT light traveled, with SIS+VEH males traveling less in the light than CNTRL+VEH males (*p*=0.029) and SIS+FLX (*p*<0.0001; Figure 2C larger panel; Bonferroni corrected). In the EPM, there was a significant effect of FLX (*F*_(1,83)_= 13.25, *p*=0.0005), with CNTRL+VEH males traveling less in EPM open arms than CNTRL+FLX (*p*=0.0161; Bonferroni corrected) and SIS+VEH males traveling less than SIS+FLX males (*p*=0.0307; Figure 2D larger panel; Bonferroni corrected). Additionally, significant SIS (*F*_(1,46)_= 7.1, *p*=0.0106) and FLX (*F*_(1,46)_= 18.92, *p*<0.0001) effects were observed in FST immobility time. Specifically, CNTRL+VEH males were more immobile than CNTRL+FLX males (*p*=0.0144), and SIS+VEH males were more immobile than SIS+FLX males (*p*=0.0026; Figure 2E large panel; Bonferroni corrected). Lastly, in the NSF, there was a significant difference in Kaplan Meier survival curves of latency to eat across groups (χ^2^_(3)_= 34.06, *p*<0.0001), with SIS+VEH males having a longer latency to feed than CNTRL+VEH (χ^2^_(1)_= 33.7, *p*<0.0001; Bonferroni corrected) and SIS+FLX (χ^2^_(1)_= 25.1, *p*<0.0001; Bonferroni corrected) males (Figure 2F, 2G). Overall in males SIS had a significant effect on OF, LDT, FST, and NSF behaviors, with FLX treatment reversing the effects of SIS stress in LDT, EPM, FST, and NSF.

Similarly, in females a 2×2 ANOVA (SISxFLX) showed a significant SIS effect (*F*_(1,91)_= 5.017, *p*=0.0276) and an interaction (*F*_(1,91)_= 4.643, *p*=0.0339) effect in the OF, with SIS+VEH females traveling less in the center than CNTRL+VEH females (*p*=0.0019; Figure 2B larger panel; Bonferroni corrected). In the LDT, a significant FLX (*F*_(1,91)_= 14.47, *p*=0.0003) effect was observed, with SIS+VEH females traveling less in the light than SIS+FLX (*p*<0.0001) and CNTRL+VEH (*p*=0.0471; Figure 2C; Bonferroni corrected) females. Within the EPM, a significant FLX (*F*_(1,91)_= 22.86, *p*<0.0001) effect was observed, with SIS+VEH females traveling less distance in the open arms than SIS+FLX females (*p*=0.0004) and CNTRL+VEH traveling less than CNTRL+FLX females (*p*=0.0039; Figure 2D; Bonferroni corrected). Furthermore, significant main effects of SIS (*F*_(1,46)_= 6.4, *p*=0.014) and FLX (*F*_(1,46)_= 24.92, *p*<0.0001) were observed in FST immobility time, with SIS+VEH females more immobile than SIS+FLX females (*p*=0.0006) and CNTRL+VEH females more immobile than CNTRL+FLX females (*p*=0.0043; Figure 2F; Bonferroni corrected). Lastly, in the NSF, there was a significant difference in Kaplan Meier survival curves of latency to eat across groups (χ^2^_(3)_= 16.41, *p*=0.0009), with SIS+VEH females having a longer NSF latency to feed than CNTRL+VEH (χ^2^_(1)_= 29.5, *p*<0.0001; Bonferroni corrected) and SIS+FLX (χ^2^_(1)_= 4.09, *p*=0.040; Figure 2F, 2H, Bonferroni corrected) females. In females, as seen in males, it appeared that SIS had an impact on OF, FST, and NSF behaviors, with FLX treatment reversing the effects of SIS stress in LDT, EPM, FST, and NSF. Taken together, these data suggest that SIS effectively induces negative valence behaviors in both adult male and adult female mice.

### Impact of Estrous Cycle on Female behavior

Since SIS impacted female behavior, we next examined the 4 stages of the estrous cycle (Proestrus, Estrus, Metestrus, Diestrus) in CNTRL and SIS females following each behavioral assay (Figure 3A). For this experiment FLX was not used. To determine SIS and estrous cycle effects on behavior, separate 2×4 ANOVAs (SIS*x*Estrous) were run for each behavioral assay. In the OF, significant estrous cycle (*F*_(3,39)_=45.16, *p*<0.001) and SIS (*F*_(1,39)_=19.13, *p*<0.0001) effects were observed. Specifically, SIS estrus females traveled less in the center than proestrus (*p*=0.0003) and metestrus (*p*=0.013) females. Additionally, SIS diestrus females had less center distance than proestrus (*p*=0.0016) and metestrus (*p*=0.0094) females (Figure 3C). Within the CNTRL group, estrus females traveled less than proestrus (*p*=0.045) or metestrus (0.0130) females, with diestrus females also traveling less than metestrus (*p*=0.023) and proestrus (*p*=0.022; larger panel Figure 3C). Similarly, in the LDT, significant estrous cycle (*F*_(3, 39)_=10.43, *p*<0.0001) and SIS (*F*_(1, 39)_=7.68, *p*=0.0085) effects were observed, with SIS estrus females traveling less in the light than proestrus (*p*=0.021) and metestrus (*p*=0.003) females. SIS diestrus females also traveled less than metestrus (*p*=0.011) females (larger panel Figure 3D). Although significant SIS effects were observed in the EPM (*F*_(1, 39)_=5.87, *p*=0.0196; Figure 3E) and FST (*F*_(1, 17)_=4.47, *p*=0.04; Figure 3F), no estrous cycle effects were found (EPM: *p*=0.849; FST: *p*=0.97). Within the OF, LDT, EPM, and FST Bonferroni corrected post-hoc comparisons did not reveal differences within estrous states (CNTRL*vs*SIS; Figures 3C-F smaller panels). Lastly, there was a significant difference in Kaplan Meier survival curves of latency to eat across estrous stages within the SIS group (χ^2^_(3)_= 11.71, *p*=0.0084), with SIS estrus females having a higher latency to feed than SIS metestrus females (χ^2^_(1)_= 15.7, *p*<0.0001; Figure 3G). Separate Kaplan Meier survival analyses showed differences in latency to eat between CNTRL and SIS females existed across estrous cycle stages (χ^2^_(8)_= 32.09, *p*<0.0001), with SIS increasing latency to eat in the proestrus (χ^2^_(1)_= 6.91, *p*=0.0085), estrus (χ^2^_(1)_= 14.93, *p*=0.0001), and diestrus (χ^2^_(1)_= 10.9, *p*=0.0001) stages (Figure 3G small panel, 3H-J). These results suggest that although the estrous cycle does increase negative valence behaviors during specific stages (mainly estrus and diestrus) the effects of SIS were still observed throughout the estrous cycle. Therefore, variability in behavior across estrous in freely cycling females does not impact the effects of SIS on behavior.

### SIS Stress Increases Exogenous Corticosterone Levels in Males and Females

We next sought to determine whether SIS led to HPA axis activation in males and females. To this end, blood was collected from mice (SIS or CNTRL) at three different time points: prior to SIS exposure, 40 minutes following a SIS cage change (during last week of SIS exposure), and 40 minutes following EPM behavior (Figure 4A). Plasma was then isolated and used to assay corticosterone levels. We first performed a 2×2×3 repeated measures ANOVA (sex x stress x time of blood collection) since group sizes were equivalent and found no main effect of sex (*p*=0.48). This result suggests that SIS may be impacting HPA axis activation similarly in males and females. Next, 2×3 repeated measures ANOVA (stress x time of blood collection) within each sex were performed. In males, we found significant effects of stress (*F*_(1, 8)_=31.48, *p*=0.0005) and time of collection (*F*_(2, 16)_=4.11, *p*=0.036) and an interaction (*F*_(2, 16)_=8.55, *p*=0.003). Specifically, SIS males had higher plasma corticosterone levels in response to a cage change (*p*=0.0054) and EPM (*p*<0.0001) than CNTRL males (Figure 4B). Significant increases in SIS male plasma corticosterone levels after cage change (*p*=0.008) and EPM exposure (*p*=0.0001) were also observed relative to levels before SIS exposure (Figure 4B). Therefore, chronic SIS exposure increased HPA axis activation in response to acute stressors in males.

**Figure 4.**
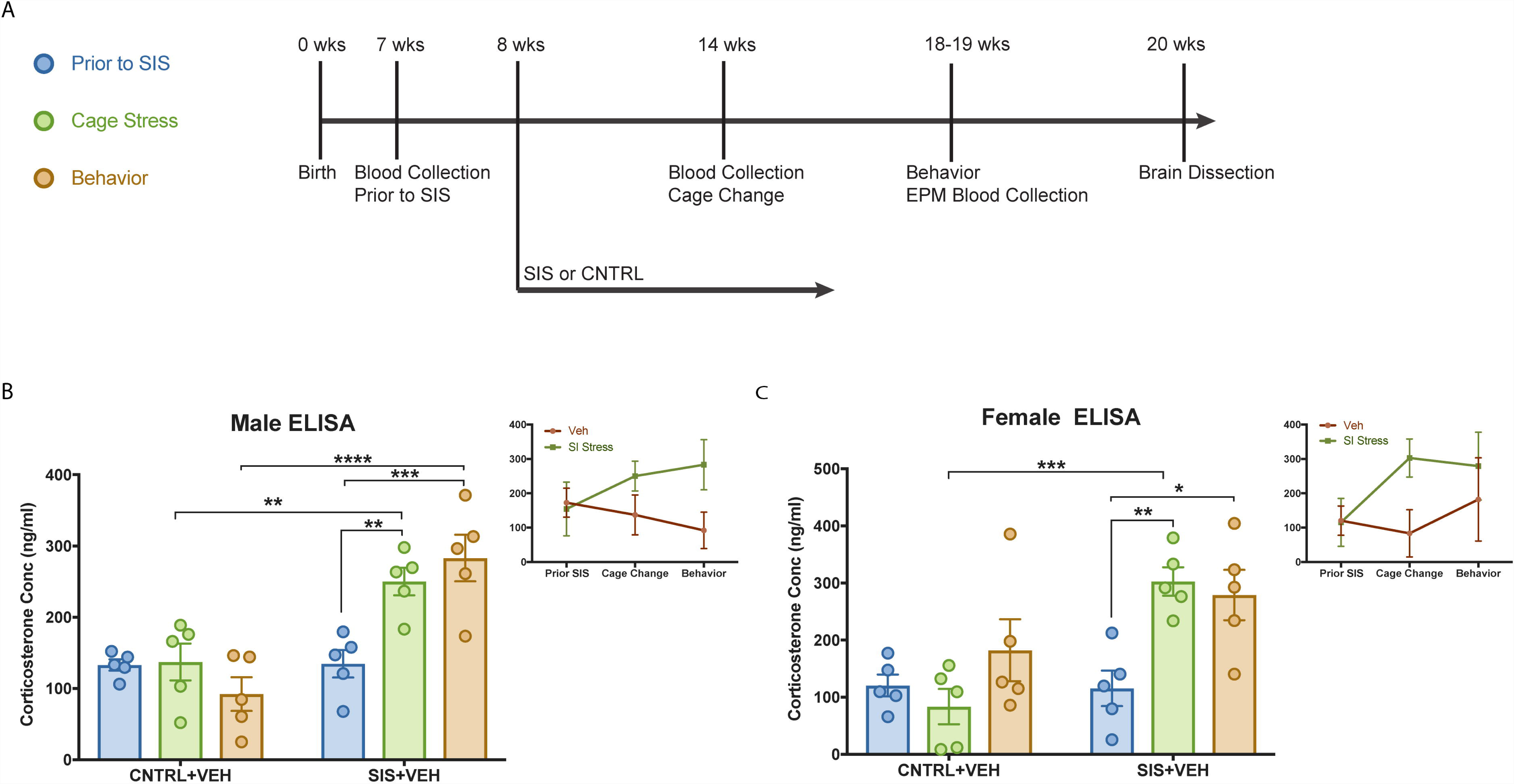
SIS Stress Increases Exogenous Corticosterone Levels in Males and Females. Timeline of experiment and legend for figures (A). Smaller panels (B-C) illustrate the changes in plasma corticosterone levels across blood collection time points. Larger panel (B) displays 2×3 ANOVAs (SIS*x*TIME) within males with SIS (*F*_(1, 8)_=31.48, *p=*0.0005) and TIME effects (*F*_(2, 16)_=4.11, *p=*0.036; **SIS+PRIOR*vs*SIS+CAGE *p*=0.008, ***SIS+PRIOR*vs*SIS+EPM *p*=0.0001) and INTERACTION (*F*_(2, 16)_=8.55, *p=*0.003, **SIS+CAGE*vs*CNTRL+CAGE *p*=0.0054; ****SIS+EPM*vs*CNTRL+EPM *p<*0.0001). In females, larger panel (C), SIS (*F*_(1, 8)_=10, *p=*0.013) and TIME effects (*F*_(2, 16)_=5.75, *p=*0.013; **SIS+PRIOR*vs*SIS+CAGE *p=*0.0037, ***SIS+PRIOR*vs*SIS+EPM *p=*0.0105) and INTERACTION (*F*_(2, 16)_=5.52, *p=*0.015, ***SIS+CAGE*vs*CNTRL+CAGE *p=*0.0007).

Separate 2×3 repeated measures ANOVAs in females revealed significant effects of stress (*F*_(1, 8)_=10, *p*=0.013) and time of collection (*F*_(2, 16)_=5.75, *p*=0.013) and an interaction (*F*_(2, 16)_=5.52, *p*=0.015). SIS females had significantly higher plasma corticosterone levels in response to a cage change than CNTRL females (*p*=0.0007; Figure 4C). Lastly, plasma corticosterone levels in SIS females were significantly higher following a cage change (*p*=0.0037) and EPM exposure (*p*=0.0105) as compared to corticosterone levels before SIS exposure (Figure 4C). Thus, similar to males, chronic SIS exposure increased the HPA axis response to acute stressors in females. Taken together, these results demonstrate that SIS leads to HPA axis activation in response to acute stressors in both males and females.

### SIS Stress and FLX Impact Adult Hippocampal Neurogenesis

In addition to behavioral and HPA axis effects, chronic stress can also affect endogenous adult hippocampal neurogenesis (44–46) and antidepressants are well-known to increase all stages of adult hippocampal neurogenesis (13, 47–49). Therefore, following behavior, mice were perfused, and brains were collected and sectioned for immunostaining. To analyze different stages of adult hippocampal neurogenesis, we immunostained for Ki67 (a proliferative marker; Figure 5A) and DCX (a newborn/immature neuron marker; Figure 5C, 6E). First, we investigated sex differences between groups (CNTRL+VEH, CNTRL+FLX, SIS+VEH, and SIS+FLX) with 2×4 ANOVAs for each hippocampal neurogenesis marker. No effects of sex were found for number of Ki67^+^ cells (*p*=0.29), DCX^+^ cells (*p*=0.45), DCX^+^ cells with tertiary dendrites (*p*=0.093), and maturation index (*p*=0.086; Figure 5B and Figure 5D, 6F-G smaller panels). Therefore, separate 2×2 ANOVAs within each sex were used to investigate SIS and FLX effects on hippocampal neurogenesis.

**Figure 5.**
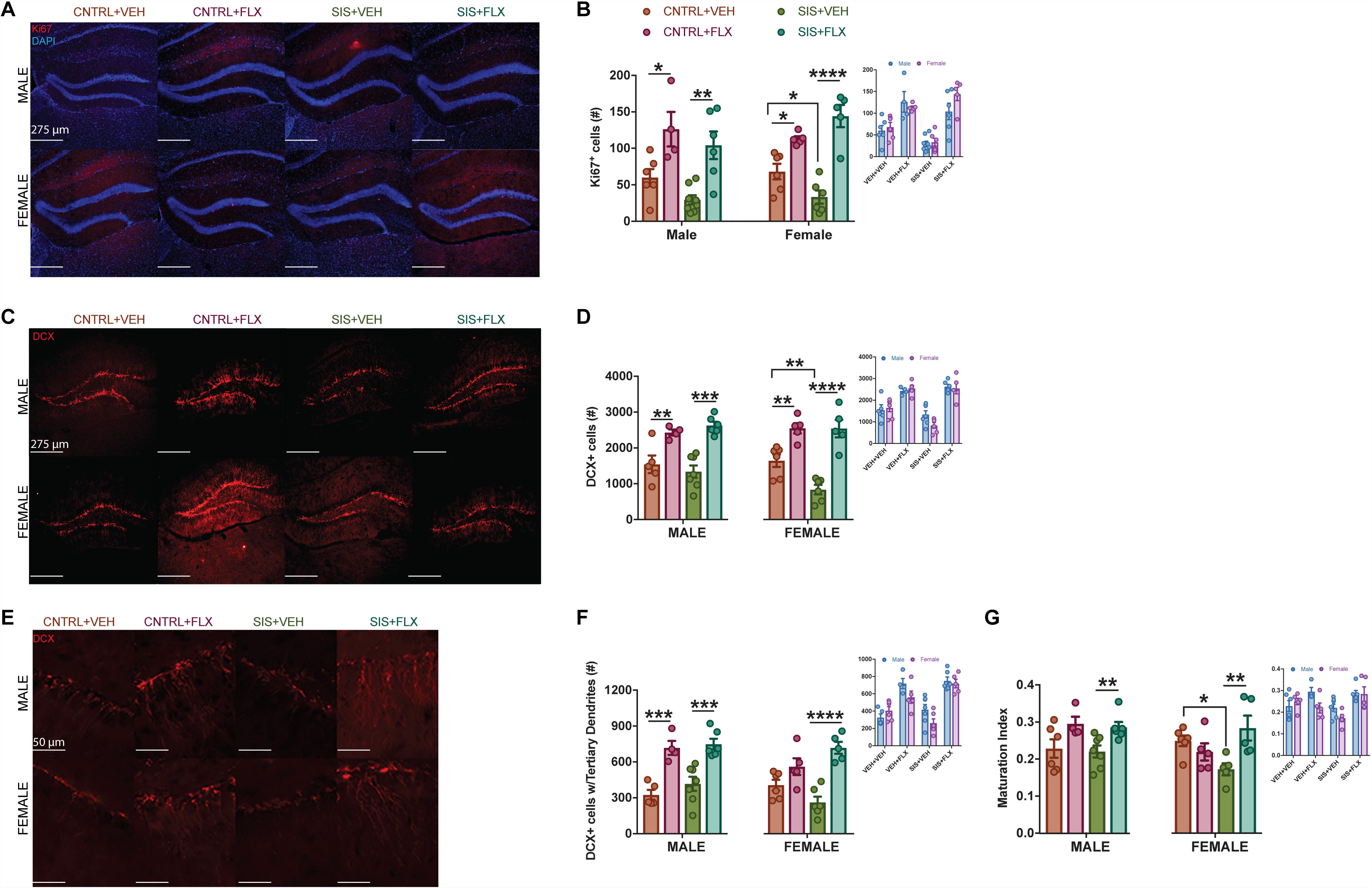
SIS Stress and FLX Effects on Adult Hippocampal Neurogenesis. Representative 10x images of Ki67 (A) and DCX (C), with 40x images used to quantify DCX^+^ cells with tertiary dendrites (E). Smaller panels (B, D, F, G) display no sex differences in adult hippocampal neurogenesis. In males 2×2 ANOVAs (SIS*x*FLX) revealed FLX effects in Ki67^+^ (*F*_(1,20)_=23.52, *p*<0.0001; *CNTRL+VEH*vs*CNTRL+FLX *p=*0.0150 and **SIS+FLX*vs*SIS+VEH *p=*0.0014), DCX^+^ (*F*_(1,18)_=30.74, *p*<0.0001; **CNTRL+FLX*vs*CNTRL+VEH *p=*0.0067 and ***SIS+FLX*vs*SIS+VEH *p=*0.0004), DCX^+^ cells with tertiary dendrites (*F*_(1,18)_=44.1, p<0.0001; ***CNTRL+FLX*vs*CNTRL+VEH *p=*0.0004 and ***SIS+FLX*vs*SIS+VEH *p=*0.0003) and maturation index (*F*_(1,19)_=36.71, *p=*0.0034; **SIS+FLX*vs*SIS+VEH *p=*0.0401). Within females 2×2 ANOVAs (SIS*x*FLX) showed main effects and interactions in Ki67 (FLX: *F*_(1,18)_=55.42, p<0.0001; INTERACTION: *F*_(1,18)_=9.968, *p=*0.0055; ****SIS+VEH*vs*SIS+FLX *p*<0.0001, *SIS+VEH*vs*CNTRL+VEH *p*=0.0475, *CNTRL+FLX*vs*CNTRL+VEH *p*=0.014), DCX (SIS: *F*_(1,18)_=5.34, *p=*0.033, FLX: *F*_(1,18)_=54.79, *p*<0.0001, INTERACTION: *F*_(1,18)_=5.12, *p*=0.036; ****SIS+VEH*vs*SIS+FLX *p*<0.0001, **SIS+VEH*vs*CNTRL+VEH *p*=0.0065, **CNTRL+FLX*vs*CNTRL+VEH *p*=0.0038), DCX^+^ cells with tertiary dendrites (INTERACTION *F*_(1,18)_=9.14, *p=*0.0073, FLX: *F*_(1,18)_=36.01, *p*<0.0001; ****SIS+VEH*vs*SIS+FLX *p*<0.0001), and maturation index (INTERACTION: *F*_(1,17)_=9.924, *p=*0.0058; **SIS+VEH*vs*SIS+FLX *p*=0.0064, *SIS+VEH*vs*CNTRL+VEH *p*=0.048).

Within males, 2×2 ANOVAs revealed a significant FLX effect (*F*_(1,20)_= 23.52, *p*<0.0001) on Ki67 expression, with FLX males having higher number of Ki67+ cells within the dentate gyrus (DG) than VEH treated males [CNTRL+VEH*vs*CNTRL+FLX (*p*=*0*.0150) and SIS+FLX*vs*SIS+VEH (*p*=*0*.0014); Figure 5A-B]. Significant FLX effects were also observed in number of DCX^+^ cells (*F*_(1,18)_= 30.74, *p*<0.0001); DCX cells maturation (*F*_(1,18)_= 44.1, *p*<0.0001); and maturation index (DCX^+^ cells with tertiary dendrites/total DCX^+^ cells; (*F*_(1,19)_= 36.71, *p*=*0*.0034), with FLX treated males having more DCX+ cells [CNTRL+FLX*vs*CNTRL+VEH (*p*=*0*.0067) and SIS+FLX*vs*SIS+VEH (*p*=*0*.0004); Figure 5D], more DCX^+^ cells with tertiary dendrites than their VEH male counterparts [CNTRL+FLX*vs*CNTRL+VEH (*p*=*0*.0004) and SIS+FLX*vs*SIS+VEH (*p*=*0*.0003); Figure 5F], and a higher maturation index (SIS+FLX*vs*SIS+VEH (*p*=*0*.0401; Figure 5G) than VEH males. Thus, in males, while FLX increased adult hippocampal neurogenesis, SIS stress did not have significant effects.

By contrast, in females separate 2×2ANOVAs revealed a significant FLX effect (*F*_(1,18)_= 55.42, *p*<0.0001) and interaction (*F*_(1,18)_= 9.968, *p*=*0*.0055) on number of DG Ki67+ cells. Specifically, SIS+VEH females had less Ki67^+^ cells than SIS+FLX (*p*<0.0001) and CNTRL+VEH (*p*=0.0475) females (Figure 5B). CNTRL+FLX females also had more Ki67^+^ cells than CNTRL+VEH (*p*=0.0144) females (Figure 5B). Significant effects of SIS (*F*_(1,18)_= 5.34, *p*=*0*.033) and FLX (*F*_(1,18)_= 54.79, *p*<0.0001), and an interaction (*F*_(1,18)_= 5.12, *p*=0.036) was also observed in number of DCX+ cells, with SIS+VEH females having less DCX^+^ cells than CNTRL+VEH (*p*=0.0065) and SIS+FLX (*p*<0.0001) females. CNTRL+FLX females also had more DCX^+^ cells than CNTRL+VEH females (*p*=0.0038; Figure 5D). Further, we observed a significant interaction (*F*_(1,18)_= 9.14, *p*=*0*.0073) and FLX effect (*F*_(1,18)_= 36.01, *p*<0.0001) on DCX maturation, with SIS+FLX females having more DCX^+^ cells with tertiary dendrites than SIS+VEH females (*p*<0.0001; Figure 5F). Lastly, a significant interaction was revealed in maturation index (*F*_(1,17)_= 9.924, *p*=0.0058), with SIS+VEH females having a lower maturation index than SIS+FLX (*p*=0.0064) and CNTRL+VEH (*p*=0.0482) females (Figure 5G). Thus, these results show that SIS stress in females does impact proliferation and differentiation.

## DISCUSSION

Here we demonstrate that the SIS paradigm effectively induces negative valence behaviors and HPA axis activation in both adult males and adult females of the widely used C57BL6/J strain. Importantly, there were no sex differences across all behavior tests following SIS exposure. These data suggest that the SIS paradigm can be used to assess the effects of chronic stress in both sexes without infringing upon ethological validity. We also found that there were sex differences following CORT administration, suggesting that the CORT paradigm to mimic chronic stress is not useful for both sexes. Importantly, we also found that while there were differences in the behavioral effects of SIS among the distinct phases of the estrous cycle, this variability did not result in any significant differences across sex when the entire cohort of females was collapsed. Therefore, SIS can be performed in adult males and freely cycling adult females without concern of estrous cycle confounds.

Our study exposes adult male and adult female C57BL/6J mice to a SIS paradigm. The most similar approaches were performed in the outbred CD-1 strain, where adolescent mice exposed to SIS displayed increased negative valence behaviors in EPM and NSF in males (37) and females (36). In these two studies, the CD-1 mice were exposed to SIS beginning at postnatal day 24, which was soon after weaning. Given that anxiety disorders can be heavily influenced by development exposure to stress (50), exposure of adolescent CD-1 mice to SIS may be similar to an early life stress paradigm. In rats, repeated resident-intruder stressful exposure of adolescent females to lactating adult females produces different patterns of effects in negative valence behaviors than exposure of adult females (51). Therefore, exposure of adolescents or adults to SIS may yield different effects. We utilized adult males and adult females from the widely used inbred C57BL/6J strain, and our data supports the notion that adult exposure to chronic stress can also impact negative valence behaviors, HPA axis activation, and adult hippocampal neurogenesis. In addition, we simultaneously ran the male and female cohorts, allowing direct comparison of potential sex differences. Furthermore, we demonstrate that subsequent FLX treatment reverses the negative valence behaviors in both males and females, lending pharmacological validity to the SIS paradigm as well. By comparing groups (CNTRL+VEH, CNTRL+FLX, SIS+VEH, and SIS+FLX) across sex, we found no sex differences of SIS exposure or FLX treatment in negative valence behaviors or adult hippocampal neurogenesis. Furthermore, there were no sex differences of SIS exposure on HPA axis activation following cage changes or EPM. Additionally, these behavioral results were validated in two separate smaller cohorts of mice, before collapsing the data across both cohorts (data not shown).

Subchronic social stress paradigms (∼15 days) involving periods of social isolation and crowding have also been assessed in rats. One study found that exposure of adolescent male rats, but not female rats, to social stress induced negative valence behaviors in EPM and HPA axis activation (52). Another study found effects of social stress in adult female rats in inducing negative valence behaviors in EPM (53). Yet another set of studies found that social stress is effective in inducing HPA axis activation in female rats, but defeat is more effective for male rats (54, 55). Finally, chronic social stress (4 weeks) combining periods of social isolation and crowding in adult female rats leads to activation of the HPA axis and a decrease in sucrose preference (56). Social stress involving periods of social isolation and crowding also leads to activation of the HPA axis but no effects in EPM or center measures of the OF in adult female CD-1 mice (57).

Given that ovarian hormones can impact stress responses and behavior (6, 58, 59) we tracked the estrous cycle throughout our behavioral experiments. We found that SIS estrus and diestrus females travel less distance in the center of the OF and in the light compartment of the LDT than SIS proestrus and metestrus females. These data are in line with findings that socially isolated female mice in the estrus and diestrus phases spend less time in the center of the open field arena than proestrus mice (6). Although behavioral differences across estrous phases were observed in both SIS and CNTRL mice, these differences did not seem to impact our results since stress effects remained when we collapsed female mice from all four stages into one group. The notion that tracking females across all stages of the estrous cycle is necessary when analyzing results is unwarranted because males are just as variable as freely cycling females (9, 10). Thus, future studies employing the SIS paradigm do not need to track estrous cycle during behavioral experiments.

Sex differences in HPA activation between males and females have been found in response to acute stressors (60–62). However, following SIS exposure, we found no sex differences in plasma corticosterone levels in response to either cage changes or EPM exposure. Both males and females exposed to SIS have an increase in plasma corticosterone levels following these acute stressors. Furthermore, prior to SIS exposure, there were no differences between males and females in plasma corticosterone. Future studies may want to assess the long-term neuroendocrine effects of SIS, by analyzing either corticosterone levels several weeks after the final exposure to unstable social environments.

Chronic stress can affect endogenous adult hippocampal neurogenesis in male rodents (44–46). Furthermore, chronic treatment with antidepressants, such as the SSRI FLX, increases all stages of adult hippocampal neurogenesis in male and female rodents (13, 47–49). To our knowledge, this is the first study to assess the effects of chronic stress and subsequent antidepressant treatment in both male and female mice simultaneously. Within females, our data shows that SIS impacts multiple stages of adult hippocampal neurogenesis, including proliferation and differentiation. Chronic FLX treatment had effects on all stages of adult neurogenesis in females. By contrast, in males, we did not see any effects of SIS on adult hippocampal neurogenesis but found effects of FLX treatment on proliferation, differentiation, and maturation. In other chronic stress paradigms, male mice administered CORT only displayed an effect on proliferation (13), while male mice exposed to CSDS had transient effects of stress on cell proliferation (63) and a reduction in total DCX^+^ cells (45). However, complete ablation of the hippocampal adult neurogenic niche using focal irradiation does not impact negative valence behaviors, suggesting that decreases in adult hippocampal neurogenesis are not sufficient or necessary to impact behavior (49). Rather, adult neurogenesis is required for the behavioral effects of antidepressant treatment. We found that FLX increased all stages of adult hippocampal neurogenesis in both males and females.

Our data suggest that the SIS paradigm is an ethologically valid approach that effectively induces chronic stress in both males and females. In contrast to another chronic stress paradigm, chronic CORT administration, we found no sex differences in the effects of SIS on negative valence behaviors and HPA axis activation. Future work is necessary to determine how long-lasting the effects of SIS are and whether SIS can be leveraged to study stress resilience and susceptibility in both males and females.

## FUNDING AND DISCLOSURE

This work was supported by NIMH (R01MH112861, BAS). The authors declare no conflicts of interest.

## ACKNOWLEDGEMENTS

The authors would like to thank Emma Diethorn and Sophie Shifman for assistance.

## AUTHOR CONTRIBUTIONS

C.N.Y. and B.A.S. conceived of experiments. C.N.Y., S.A.A., L.B., and A.G. performed the experiments. C.N.Y., M.M.G., and B.A.S. analyzed the data and made the figures. C.N.Y., M.M.G., and B.A.S. wrote the manuscript.

## REFERENCES

1. Beery, A. K., & Zucker, I. (2011). Sex bias in neuroscience and biomedical research. Neurosci Biobehav Rev, 35(3), 565–572.

2. Autry AE, Adachi M, Cheng P, Monteggia LM (2009). Genderspecific impact of brain-derived neurotrophic factor signaling on stress-induced depression-like behavior. Biol Psychiatry 66: 84–90.

3. Trainor, B.C. (2011). Stress responses and the mesolimbic dopamine system: social contexts and sex differences. Horm Behav 60: 457–469.

4. Shansky RM (2015). Sex differences in PTSD resilience and susceptibility: challenges for animal models of fear learning. Neurobiol Stress 1: 60–65.

5. Becker, J. B., Arnold, A. P., Berkley, K. J., Blaustein, J. D., Eckel, L. A., Hampson, E., … & Taylor, J. (2005). Strategies and methods for research on sex differences in brain and behavior. Endocrinology, 146(4), 1650–1673.

6. Palanza, P., Gioiosa, L., & Parmigiani, S. (2001). Social stress in mice: gender differences and effects of estrous cycle and social dominance. Physiol Behav, 73(3), 411–420.

7. Shansky, R. M., & Woolley, C. S. (2016). Considering sex as a biological variable will be valuable for neuroscience research. J Neurosci, 36(47), 11817–11822.

8. Yohn, C. N., Shifman, S., Garino, A., Diethorn, E., Bokka, L., Ashamalla, S. A., & Samuels, B. A. (2018). Fluoxetine effects on behavior and adult hippocampal neurogenesis in female C57BL/6J mice across the estrous cycle. bioRxiv, 368449.

9. Becker, J. B., Prendergast, B. J., & Liang, J. W. (2016). Female rats are not more variable than male rats: a meta-analysis of neuroscience studies. Biol Sex Differ, 7(1), 34.

10. Prendergast, B. J., Onishi, K. G., & Zucker, I. (2014). Female mice liberated for inclusion in neuroscience and biomedical research. Neurosci Biobehav Rev, 40, 1–5.

11. Shansky, R. M. (2018). Sex differences in behavioral strategies: avoiding interpretational pitfalls. Current opinion in neurobiology, 49, 95–98.

12. Blanchard, D. C., Griebel, G., & Blanchard, R. J. (1995). Gender bias in the preclinical psychopharmacology of anxiety: male models for (predominantly) female disorders. J Psychopharmacol, 9(2), 79–82.

13. David, D. J., Samuels, B. A., Rainer, Q., Wang, J. W., Marsteller, D., Mendez, I., … & Artymyshyn, R. P. (2009). Neurogenesis-dependent and-independent effects of fluoxetine in an animal model of anxiety/depression. Neuron, 62(4), 479–493.

14. Golden, S. A., Covington III, H. E., Berton, O., & Russo, S. J. (2011). A standardized protocol for repeated social defeat stress in mice. Nat Protoc, 6(8), 1183.

15. Russo, S. J., & Nestler, E. J. (2013). The brain reward circuitry in mood disorders. Nat Rev Neurosci, 14(9), 609.

16. Mineur, Y. S., Belzung, C., & Crusio, W. E. (2006). Effects of unpredictable chronic mild stress on anxiety and depression-like behavior in mice. Behav Brain Res, 175(1), 43–50.

17. Piantadosi, S. C., French, B. J., Poe, M. M., Timić, T., Marković, B. D., Pabba, M., … & Cook, J. M. (2016). Sex-dependent anti-stress effect of an α5 subunit containing GABAA receptor positive allosteric modulator. Front Pharmacol, 7, 446.

18. Guilloux, J. P., Seney, M., Edgar, N., & Sibille, E. (2011). Integrated behavioral z-scoring increases the sensitivity and reliability of behavioral phenotyping in mice: relevance to emotionality and sex. J Neurosci Meth, 197(1), 21–31.

19. Zhao, Z., Zhang, L., Guo, X. D., Cao, L. L., Xue, T. F., Zhao, X. J., … & Sun, X. L. (2017). Rosiglitazone exerts an anti-depressive effect in unpredictable chronic mild-stress-induced depressive mice by maintaining essential neuron autophagy and inhibiting excessive astrocytic apoptosis. Front Mol Neurosci, 10, 293.

20. Belzung, C., & Lemoine, M. (2011). Criteria of validity for animal models of psychiatric disorders: focus on anxiety disorders and depression. Biology of mood & anxiety disorders, 1(1), 9.

21. Farooq, R. K., Isingrini, E., Tanti, A., Le Guisquet, A. M., Arlicot, N., Minier, F., … & Camus, V. (2012). Is unpredictable chronic mild stress (UCMS) a reliable model to study depression-induced neuroinflammation?. Behav Brain Res, 231(1), 130–137.

22. Willner, P. (1997). Validity, reliability and utility of the chronic mild stress model of depression: a 10-year review and evaluation. Psychopharmacology, 134(4), 319–329.

23. Krishnan, V., Han, M. H., Graham, D. L., Berton, O., Renthal, W., Russo, S. J., … & Ghose, S. (2007). Molecular adaptations underlying susceptibility and resistance to social defeat in brain reward regions. Cell, 131(2), 391–404

24. Keeney, A., Jessop, D. S., Harbuz, M. S., Marsden, C. A., Hogg, S., & Blackburn-Munro, R. E. (2006). Differential effects of acute and chronic social defeat stress on hypothalamic-pituitary-adrenal axis function and hippocampal serotonin release in mice. J Neuroendocrinol, 18(5), 330–338.

25. Greenberg, G. D., Laman-Maharg, A., Campi, K. L., Voigt, H., Orr, V. N., Schaal, L., & Trainor, B. C. (2014). Sex differences in stress-induced social withdrawal: role of brain derived neurotrophic factor in the bed nucleus of the stria terminalis. Front Behav Neurosci, 7, 223.

26. Harris, A. Z., Atsak, P., Bretton, Z. H., Holt, E. S., Alam, R., Morton, M. P., … & Gordon, J. A. (2018). A novel method for chronic social defeat stress in female mice. Neuropsychopharmacol, 43(6), 1276.

27. Takahashi, A., Chung, J. R., Zhang, S., Zhang, H., Grossman, Y., Aleyasin, H., … & Hodes, G. E. (2017). Establishment of a repeated social defeat stress model in female mice. Sci Rep, 7(1), 12838.

28. Herman, J. P., McKlveen, J. M., Ghosal, S., Kopp, B., Wulsin, A., Makinson, R., … & Myers, B. (2016). Regulation of the hypothalamic-pituitary-adrenocortical stress response. Compr Physiol, 6(2), 603

29. Kitay, J. I. (1963). Pituitary-adrenal function in the rat after gonadectomy and gonadal hormone replacement. Endocrinology, 73(2), 253–260.

30. Wardenaar, K. J., Vreeburg, S. A., van Veen, T., Giltay, E. J., Veen, G., Penninx, B. W., & Zitman, F. G. (2011). Dimensions of depression and anxiety and the hypothalamo-pituitary-adrenal axis. Biol Psychiatry, 69(4), 366–373.

31. Gourley, S. L., Wu, F. J., & Taylor, J. R. (2008). Corticosterone regulates pERK1/2 map kinase in a chronic depression model. Ann NY Acad Sci, 1148(1), 509–514.

32. Gourley, S. L., & Taylor, J. R. (2011). Induction of persistent depressive-like behavior by corticosterone. In Mood and Anxiety Related Phenotypes in Mice (pp. 251–265). Humana Press.

33. Brummelte, S., & Galea, L. A. (2010). Chronic corticosterone during pregnancy and postpartum affects maternal care, cell proliferation and depressive-like behavior in the dam. Horm Behav, 58(5), 769–779.

34. Brummelte, S., Pawluski, J. L., & Galea, L. A. (2006). High post-partum levels of corticosterone given to dams influence postnatal hippocampal cell proliferation and behavior of offspring: a model of post-partum stress and possible depression. Horm Behav, 50(3), 370–382.

35. Mekiri, M., Gardier, A. M., David, D. J., & Guilloux, J. P. (2017). Chronic corticosterone administration effects on behavioral emotionality in female c57bl6 mice. Exp Clin Psychopharmacol, 25(2), 94.

36. Schmidt, M. V., Scharf, S. H., Liebl, C., Harbich, D., Mayer, B., Holsboer, F., & Müller, M. B. (2010). A novel chronic social stress paradigm in female mice. Horm Behav, 57(4-5), 415–420.

37. Sterlemann, V., Ganea, K., Liebl, C., Harbich, D., Alam, S., Holsboer, F., … & Schmidt, M. V. (2008). Long-term behavioral and neuroendocrine alterations following chronic social stress in mice: implications for stress-related disorders. Horm Behav, 53(2), 386–394

38. Schmidt, M. V., Sterlemann, V., Ganea, K., Liebl, C., Alam, S., Harbich, D., … & Müller, M. B. (2007). Persistent neuroendocrine and behavioral effects of a novel, etiologically relevant mouse paradigm for chronic social stress during adolescence. Psychoneuroendocrinol, 32(5), 417–429.

39. Bartolomucci, A., Pederzani, T., Sacerdote, P., Panerai, A. E., Parmigiani, S., & Palanza, P. (2004). Behavioral and physiological characterization of male mice under chronic psychosocial stress. Psychoneuroendocrinol, 29(7), 899–910.

40. McLean, A. C., Valenzuela, N., Fai, S., & Bennett, S. A. (2012). Performing vaginal lavage, crystal violet staining, and vaginal cytological evaluation for mouse estrous cycle staging identification. Journal of visualized experiments: JoVE, (67).

41. Byers, S. L., Wiles, M. V., Dunn, S. L., & Taft, R. A. (2012). Mouse estrous cycle identification tool and images. PloS ONE, 7(4), e35538.

42. Felicio, L. S., Nelson, J. F., & Finch, C. E. (1984). Longitudinal studies of estrous cyclicity in aging C57BL/6J mice: II. Cessation of cyclicity and the duration of persistent vaginal cornification. Biol Reprod, 31(3), 446–453.

43. Franklin, K. B., & Paxinos, G. (2008). The mouse brain in stereotaxic coordinates, compact. The coronal plates and diagrams. Amsterdam: Elsevier Academic Press.

44. Czéh, B., Welt, T., Fischer, A. K., Erhardt, A., Schmitt, W., Müller, M. B., … & Keck, M. E. (2002). Chronic psychosocial stress and concomitant repetitive transcranial magnetic stimulation: effects on stress hormone levels and adult hippocampal neurogenesis. Biol Psychiatry, 52(11), 1057–1065.

45. Van Bokhoven, P., Oomen, C. A., Hoogendijk, W. J. G., Smit, A. B., Lucassen, P. J., & Spijker, S. (2011). Reduction in hippocampal neurogenesis after social defeat is longlasting and responsive to late antidepressant treatment. Eur J Neurosci, 33(10), 1833–1840.

46. Pham, K., Nacher, J., Hof, P. R., & McEwen, B. S. (2003). Repeated restraint stress suppresses neurogenesis and induces biphasic PSA-NCAM expression in the adult rat dentate gyrus. Eur J Neurosci, 17(4), 879–886.

47. Malberg, J. E., Eisch, A. J., Nestler, E. J., & Duman, R. S. (2000). Chronic antidepressant treatment increases neurogenesis in adult rat hippocampus. J Neurosci, 20(24), 9104–9110.

48. Hill, A. S., Sahay, A., & Hen, R. (2015). Increasing adult hippocampal neurogenesis is sufficient to reduce anxiety and depression-like behaviors. Neuropsychopharmacol, 40(10), 2368.

49. Santarelli, L., Saxe, M., Gross, C., Surget, A., Battaglia, F., Dulawa, S., … & Belzung, C. (2003). Requirement of hippocampal neurogenesis for the behavioral effects of antidepressants. Science, 301(5634), 805–809.

50. Leonardo, E. D., & Hen, R. (2008) Anxiety and developmental disorders. Neuropsychopharm, 33 (1), 134.

51. Ver Hoeve, E. S., Kelly, G., Luz, S., Ghanshani, S., & Bhatnagar, S. (2013). Short-term and long-term effects of repeated social defeat during adolescence or adulthood in female rats. Neuroscience, 249, 63–73.

52. Roeckner, A. R., Bowling, A., & Butler, T. R. (2017). Chronic social instability increases anxiety-like behavior and ethanol preference in male Long Evans rats. Physiol Behav, 173, 179–187.

53. Haller, J., Leveleki, C., Baranyi, J., Mikics, E., & Bakos, N. (2003). Stress, social avoidance and anxiolytics: a potential model of stress-induced anxiety. Behav Pharmacol, 14(5), 439–446.

54. Haller, J., Fuchs, E., Halász, J., & Makara, G. B. (1999). Defeat is a major stressor in males while social instability is stressful mainly in females: towards the development of a social stress model in female rats. Brain Res Bull, 50(1), 33–39.

55. Nowacka, M. M., Paul-Samojedny, M., Bielecka, A. M., Plewka, D., Czekaj, P., & Obuchowicz, E. (2015). LPS reduces BDNF and VEGF expression in the structures of the HPA axis of chronic social stressed female rats. Neuropeptides, 54, 17–27.

56. Herzog, C. J., Czéh, B., Corbach, S., Wuttke, W., Schulte-Herbrüggen, O., Hellweg, R., … & Fuchs, E. (2009). Chronic social instability stress in female rats: a potential animal model for female depression. Neuroscience, 159(3), 982–992.

57. Jarcho, M. R., Massner, K. J., Eggert, A. R., & Wichelt, E. L. (2016). Behavioral and physiological response to onset and termination of social instability in female mice. Horm Behav, 78, 135–140.

58. Sisk, C. L., & Zehr, J. L. (2005). Pubertal hormones organize the adolescent brain and behavior. Front Neuroendocrinol, 26(3-4), 163–174

59. Wood, G. E., Beylin, A. V., & Shors, T. J. (2001). The contribution of adrenal and reproductive hormones to the opposing effects of stress on trace conditioning males versus females. Behav Neurosci, 115(1), 175.

60. Kirschbaum, C., Kudielka, B. M., Gaab, J., Schommer, N. C., & Hellhammer, D. H. (1999). Impact of gender, menstrual cycle phase, and oral contraceptives on the activity of the hypothalamus-pituitary-adrenal axis. Psychosom Med, 61(2), 154–162.

61. Kirschbaum, C., Wüst, S., & Hellhammer, D. (1992). Consistent sex differences in cortisol responses to psychological stress. Psychosom Med, 54(6), 648–657.

62. Kudielka, B. M., & Kirschbaum, C. (2005). Sex differences in HPA axis responses to stress: a review. Biol Psychology, 69(1), 113–132.

63. Lagace, D. C., Donovan, M. H., DeCarolis, N. A., Farnbauch, L. A., Malhotra, S., Berton, O., … & Eisch, A. J. (2010). Adult hippocampal neurogenesis is functionally important for stress-induced social avoidance. Proceedings of the National Academy of Sciences, 200910072.

